# Developmental axon diameter growth of central nervous system axons does not depend on ensheathment or myelination by oligodendrocytes

**DOI:** 10.1101/2025.01.10.632348

**Authors:** Jenea M Bin, Katie Emberley, Tobias J Buscham, Maria A Eichel-Vogel, Ryan A Doan, Anna M Steyer, Matthew F Nolan, Wiebke Möbius, Kelly R Monk, Hauke B Werner, Ben Emery, David A Lyons

## Abstract

Myelination facilitates the rapid conduction of action potentials along axons. In the central nervous system (CNS), myelinated axons vary over 100-fold in diameter, with conduction speed scaling linearly with increasing diameter. Axon diameter and myelination are closely interlinked, with axon diameter exerting a strong influence on myelination. Conversely, myelinating Schwann cells in the peripheral nervous system can both positively and negatively affect axon diameter. However, whether axon diameter is regulated by CNS oligodendrocytes is less clear. Here, we investigated CNS axon diameter growth in the absence of myelin using mouse (*Mbp^shi/shi^* and M*yrf* conditional knockout) and zebrafish (*olig2* morpholino) models. We find that neither the ensheathment of axons, nor the formation of compact myelin are required for CNS axons to achieve appropriate and diverse diameters. This indicates that developmental CNS axon diameter growth is independent of myelination, and shows that myelinating cells of CNS and PNS differentially influence axonal morphology.

## INTRODUCTION

In the vertebrate nervous system, two principal mechanisms have evolved to facilitate the rapid conduction of action potentials along axons: increasing axonal diameter and the myelination of axons. While myelination provides a means to increase speed without increasing axon diameter, axon diameters between myelinated CNS axons still vary over 100-fold ^1, 2, 3^. In general, conduction speeds along myelinated axons scale linearly with axon diameter ^4, 5^; thus, diversity in myelinated axon diameter provides the nervous system with a huge capacity to control the speed of axon potential propagation and contributes to the precise timings required for nervous system function ^6, 7, 8, 9^. Despite this importance, there remains large gaps in our knowledge as to how myelinated axon diameter is regulated.

The health and function of myelinated axons is dependent on a multitude of bidirectional interactions. Myelin affects the underlying axon by organizing its excitable domains ^10^ and providing support via the transfer of metabolites and extracellular vesicles ^11, 12^. Conversely, signals from the axon, including axon diameter, influence which axons get myelinated, as well as the length and thickness of myelin sheaths ^13, 14, 15, 16, 17, 18^. However, comparatively little is known about how myelin affects the growth of axons in diameter. In fact, myelination often occurs before axons have finished growing in diameter and thus has the potential to both promote or restrict diameter growth. For example, the growth of axons to large diameters may rely on trophic or metabolic support from myelin. Alternatively, the presence of myelin may signal to axons a reduced need for diameter growth that would otherwise be required to increase conduction speeds, or it may physically restrict the growth of axons in diameter.

Indeed, in the peripheral nervous system (PNS) there is evidence that myelin can both positively and negatively affect axon diameter. Schwann cell manipulations that result in fewer axons becoming myelinated or in thinner myelin sheaths are associated with reduced axon diameters ^19, 20, 21, 22^. Similarly, deficiency of myelin-associated glycoprotein (MAG) leads to smaller diameter myelinated axons ^23, 24^, suggesting roles for myelinating Schwann cells in promoting or maintaining diameter growth. In contrast, loss of the myelin protein chemokine-like factor-like MARVEL-transmembrane domain-containing family member-6 (CMTM6) leads to axon expansion in the absence of any notable myelin pathologies, indicating that signals from myelinating Schwann cell also restrict axon diameter ^23^.

Fewer studies have explored the role of myelinating oligodendrocytes in regulating axon diameter within the CNS, where available space for diameter growth is constrained by the skull and vertebral column. However, there is evidence that the influence of myelinating glia in the CNS may be different than in the PNS, which is perhaps not surprising given that the developmental origin, structure (e.g. myelin periodicity, number of myelin sheathes per myelinating cell) and protein composition of myelin formed by oligodendrocytes and Schwann cells are not the same ^25, 26, 27^. For example, in the CNS loss of MAG was found to result in larger, not smaller, diameter axons ^28^, and CMTM6 is not expressed by oligodendrocytes ^29^, indicating that myelin may not regulate diameter in the same way. Furthermore, while it is often interpreted that myelin regulates the growth of CNS axons in diameter, disruption of myelin composition and morphology often leads to complex pathologies. It remains unclear whether changes in diameter observed in myelin mutants ^28, 30^, or following demyelination ^31^, in the CNS represent a role for myelin in promoting or restricting normal diameter growth, or whether axon diameter is changed as a consequence of abnormal myelin. Here, we performed an in-depth analysis of axon diameter growth in the absence of CNS myelin, oligodendrocyte ensheathment, or oligodendrocytes in mouse and zebrafish models and find that developmental growth of axons in diameter is unaffected. Together these findings show that axons in the CNS do not rely on ensheathment or myelination by oligodendrocytes to grow to appropriate and diverse diameters and highlight a difference in the role of myelin in regulating developmental axon diameter growth between the CNS and PNS.

## RESULTS

### Absence of compact myelin does not impact axon diameter in the Shiverer mouse optic nerve

To address the role of myelin on axon diameter in the CNS, we first used the well-established *Mbp^shi/shi^* (*shiverer*) mouse model ^32^. The *shiverer* (*shi*) allele is characterized by a large deletion within the gene encoding myelin basic protein (MBP)^33, 34^ and *Mbp^shi/shi^* mice display no detectable *Mbp* mRNA or protein ^35, 36, 37^. This results in a severe CNS dysmyelination, while myelin in the PNS remains largely normal ^38, 39^. Oligodendrocytes in *Mbp^shi/shi^* mice are able to extend processes which contact axons, and a subset of axons are wrapped/ensheathed by oligodendrocyte membrane; however, these wraps do not become properly compacted into mature myelin sheaths ^40, 41, 42^. Previous studies examining axon diameter within the optic nerve of *Mbp^shi/shi^* mice have reported conflicting findings regarding how the absence of compact myelin influences the growth of axons in diameter. Kirkpatrick et al ^43^ reported an increased proportion of smaller diameter axons suggesting that the formation of compact myelin was required for axons to reach their appropriate diameters. However, Sanchez et al ^44^ reported oligodendrocyte ensheathment, in the absence of compact myelin was sufficient for axons to reach their appropriate diameters in *Mbp^shi/shi^* mice.

We revisited the *Mbp^shi/shi^* model using an increased sample number, and ensuring equal and unbiased sampling of axons across entire optic nerve cross sections of *Mbp^shi/shi^* mice and their wild-type littermates (see Methods). The optic nerve, which is composed of retinal ganglion cell axons, starts to be myelinated around postnatal day (P)5 and is almost completely myelinated (∼95% of axons) by P75 in wild-type mice ^45^. We assessed axon diameters in transmission electron micrograph images of optic nerve cross sections obtained from P75 *Mbp^shi/shi^* mice and their wild-type littermates, when myelination is well-established (Figure 1 A,B). There was no difference in the median axon diameter in the *Mbp^shi/shi^* mice compared to wild-type littermates at this age (Figure 1 C, Supplemental Figure 1). We also did not observe any significant difference in the distribution of axon diameters observed within the optic nerve of *Mbp^shi/shi^* mice (Figure 1 D,E, Supplemental Figure 1), with both wild-type and *Mbp^shi/shi^* axons exhibiting a roughly 10-fold range in diameter. Together, these data indicate that optic nerve axons grow to their normal and diverse diameters in the absence of compact myelin.

**Figure 1:**
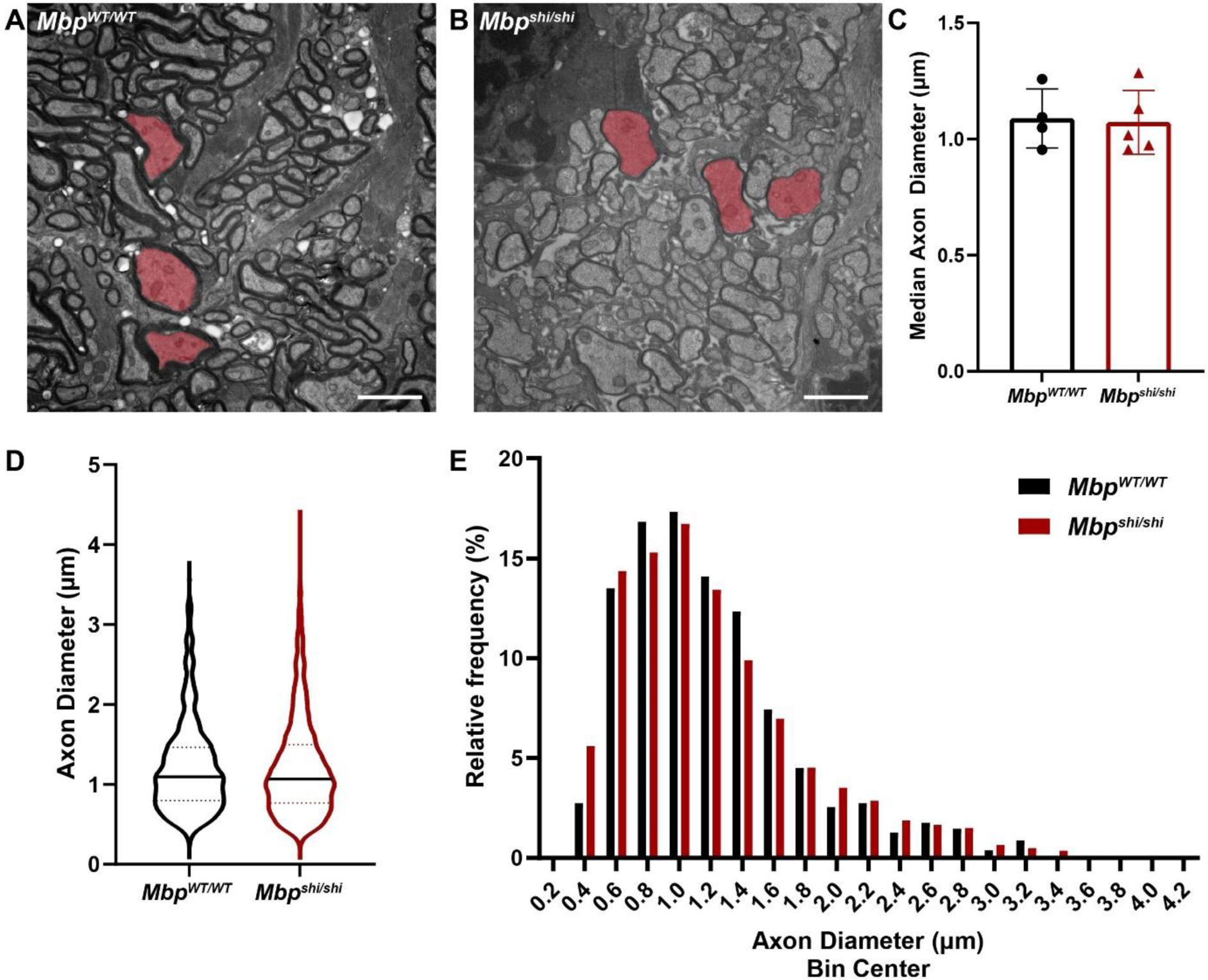
Normal axon diameters in optic nerves of *Mbp^shi/shi^* mice in the absence of compact myelin. (A-B) Representative transmission electron micrograph images from cross sections of the optic nerve at P75 for wild-type (*Mbp^WT/WT^)* (A) and *Mbp^shi/shi^* (B) mice. Shaded red axons highlight large diameter axons present in both genotypes. (C) Median axon diameter per mouse (mean +/− standard deviation for *Mbp^WT/WT^* 1.089 +/− 0.1270, n=4; *Mbp^shi/shi^* 1.072 +/− 0.1371, n=5, two-tailed unpaired t-test, p=0.8597). (D) Violin plots showing the distribution of axon diameters measured (*Mbp^WT/WT^* n=4 mice, 219-291 axons per animal; *Mbp^shi/shi^* n=5 mice, 259-290 axons per animal). Solid horizontal line marks the median and horizontal dotted lines mark the quartiles. See Supplemental Figure 1 for distributions per animal and statistical analysis. (E) Relative frequency distribution of axons in (D) divided in 0.2 µm bins. Scale bars = 2 µm.

### Early axon diameter growth is unaffected in the absence of oligodendrocyte ensheathment

While the *Mbp^shi/shi^* mice lack compact myelin, oligodendrocytes processes still contact and ensheathe a subset of axons and may therefore influence axon diameter via myelin-independent signaling mechanisms and/or trophic support. Therefore, we next asked how axon diameter is influenced by the absence of both myelination and oligodendrocyte ensheathment by conditionally disrupting expression of myelin regulatory factor (MYRF) in oligodendrocytes using *Myrf^fl/fl^; Olig2^wt/cre^* mice ^46^. MYRF is a transcription factor that is necessary for CNS myelination. In the absence of MYRF, oligodendrocyte precursor cells fail to differentiate into mature oligodendrocytes and ensheathe axons ^46^. We examined axon diameter in the optic nerve of *Myrf^fl/fl^; Olig2^wt/cre^* mice (*Myrf* cKO) and *Myrf^fl/+^; Olig2^wt/cre^* (control) littermates at P14. This is the latest timepoint we could assess, as mutant animals develop severe tremors and ataxia and die during the third postnatal week ^46^. We have previously shown that by this timepoint, approximately 50% of retinal ganglion cell axons are ensheathed or myelinated in controls, versus less than 0.1% of axons being ensheathed in conditional mutants ^46^.

In line with the greatly reduced number of oligodendrocytes ^46^, we found that at P14 the cross-sectional area of the *Myrf* cKO optic nerves was roughly 43% smaller than that of control optic nerves (Figure 2 A,D). By transmission electron microscopy, there was a notable absence of oligodendrocytes processes and myelin, with axons packed 1.8x more densely than in controls (Figure 2 B,C,E). Quantification of axon number revealed a similar number in control and *Myrf* cKO mice, suggesting axon loss did not contribute to the smaller nerve size (Figure 2 F). Sampling of axon diameters throughout the optic nerve indicated that there were no differences in the median or distribution of axon diameters in *Myrf* cKO mice (Figure 2 G-I, Supplemental Figure 2), with the same proportion of mutant axons reaching the diameter of myelinated axons in control mice (Figure J). This indicates that the population of larger diameter myelinated axons in wild-type mice are not larger due to their being myelinated, but rather that larger axons are those that are first selected for myelination. While we can not exclude later roles for oligodendrocyte ensheathment in regulating or maintaining axon diameter, these results indicate that the early developmental and differential diameter growth of retinal ganglion cell axons in the optic nerve is not dependent on ensheathment or myelination by oligodendrocytes.

**Figure 2:**
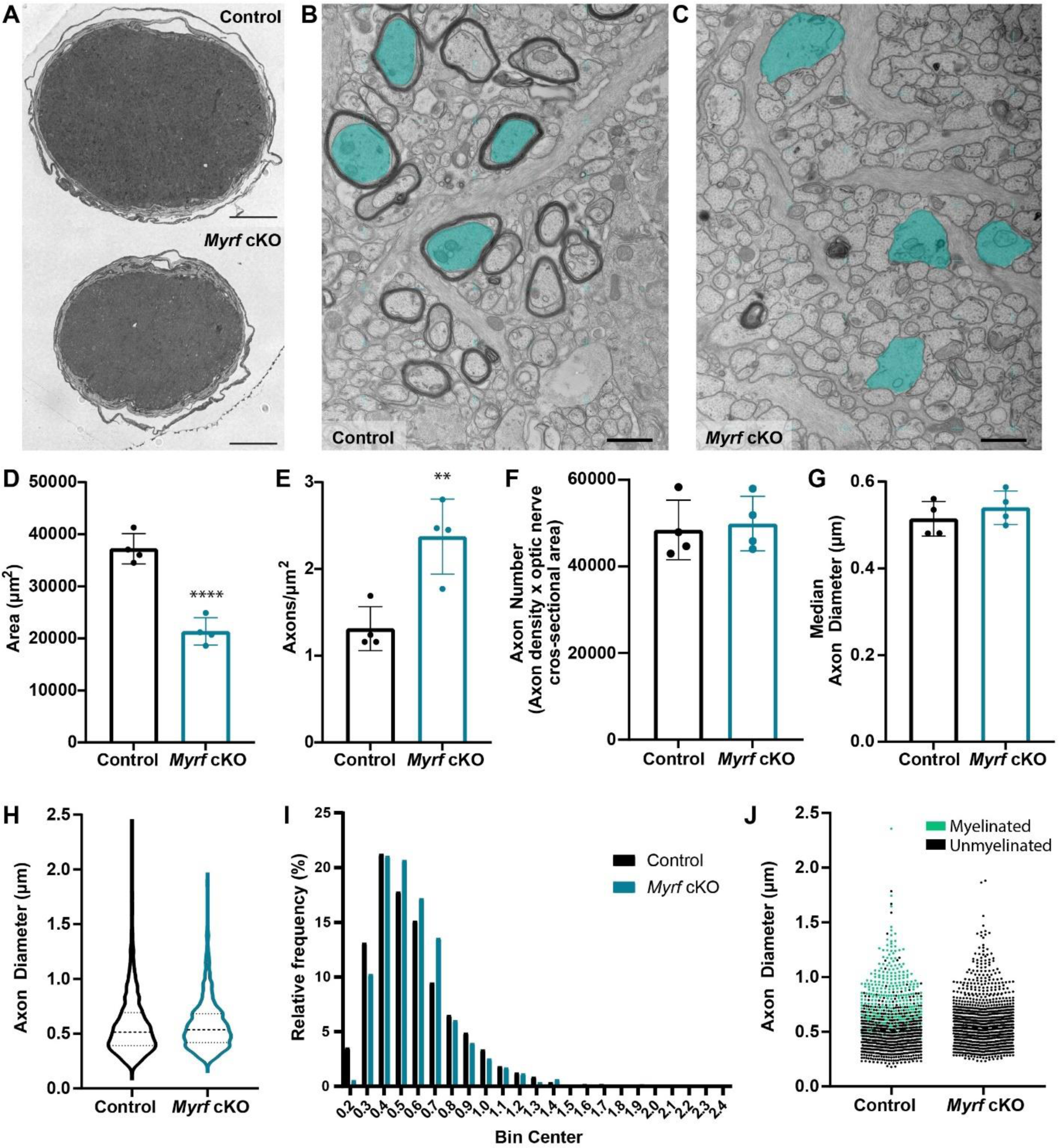
Normal axon diameters in optic nerves of *Myrf* cKOs in the absence of oligodendrocyte ensheathment and myelination. (A) Representative overview images of the optic nerve cut in cross section for control and *Myrf* cKO. Scale bars = 50 µm. (B-C) Representative transmission electron micrographs from cross sections of the optic nerve at P14 for control (B) and *Myrf* cKO (C) mice. Shaded blue axons highlight large diameter axons present in both genotypes. Scale bars = 1 µm. (D) Cross-sectional area of the optic nerve (mean +/− standard deviation for control 37220 +/− 2914, n=4; *Myrf* cKO 21331 +/− 2618, n=4, two-tailed unpaired t-test, ****p=0.0002). (E) Axon density in the optic nerve (mean +/− standard deviation for control 1.31 +/− 0.25, n=4; *Myrf* cKO 2.73+/− 0.43, n=4, two-tailed unpaired t-test, **p=0.0055). (F) Number of axons in the optic nerve estimated using the axon density and optic nerve cross-sectional area measurements (mean +/− standard deviation for control 48467 +/− 6874, n=4; *Myrf* cKO 49937+/− 6302, n=4, two-tailed unpaired t-test, p=0.7633). (G) Median axon diameter in the optic nerve per mouse (mean +/− standard deviation for control 0.514 +/− 0.040, n=4; *Myrf* cKO 0.5398+/− 0.039, n=4, two-tailed unpaired t-test, p=0.3911). (H) Violin plots showing the distribution of diameters for all axons measured (control n=4 mice, 214-301 axons per animal; *Myrf* cKO n=4 mice, 260-290 axons per animal). Solid horizontal line marks the median and horizontal dotted lines mark the quartiles. See Supplemental Figure 2 for distributions per animal and statistical analysis. (I) Relative frequency distribution of axons in (H) divided in 0.2 µm bins. (J) Axon diameters for all axons measured color coded based on whether the axon is myelinated (green) or unmyelinated (black).

### Axon diameter growth in the spinal cord of zebrafish is unaffected by the absence of oligodendrocytes and myelin

As an additional model to examine axon diameter growth in the absence of oligodendrocytes, we turned to the zebrafish spinal cord, which also offered the additional advantage of being able to follow diameter growth of specific identifiable neurons over time. Myelination in the zebrafish spinal cord begins around 2.5 days post-fertilization (dpf) ^47^, with robust myelination observed within both the dorsal and ventral tracts by 5 dpf (Figure 3 A-D). At this timepoint, axon diameters already vary by over 40-fold between neurons ^48^, with the largest of these axons belonging to a bilateral pair of reticulospinal neurons known as the Mauthner neurons ^49^. We have previously characterized the diameter growth of the Mauthner axon in relation to its myelination using time course live-imaging and shown that within the first three days of its myelination, the axon grows 3-4x in diameter ^48^. To determine if this growth in diameter is regulated by oligodendrocytes we disrupted function of Olig2, a transcription factor required for the specification of oligodendrocyte precursor cells from neural progenitors within the spinal cord ^50, 51, 52, 53^. To do this, we used an established *olig2* morpholino (MO)-based knockdown protocol ^54^ to block translation of Olig2 protein.

**Figure 3:**
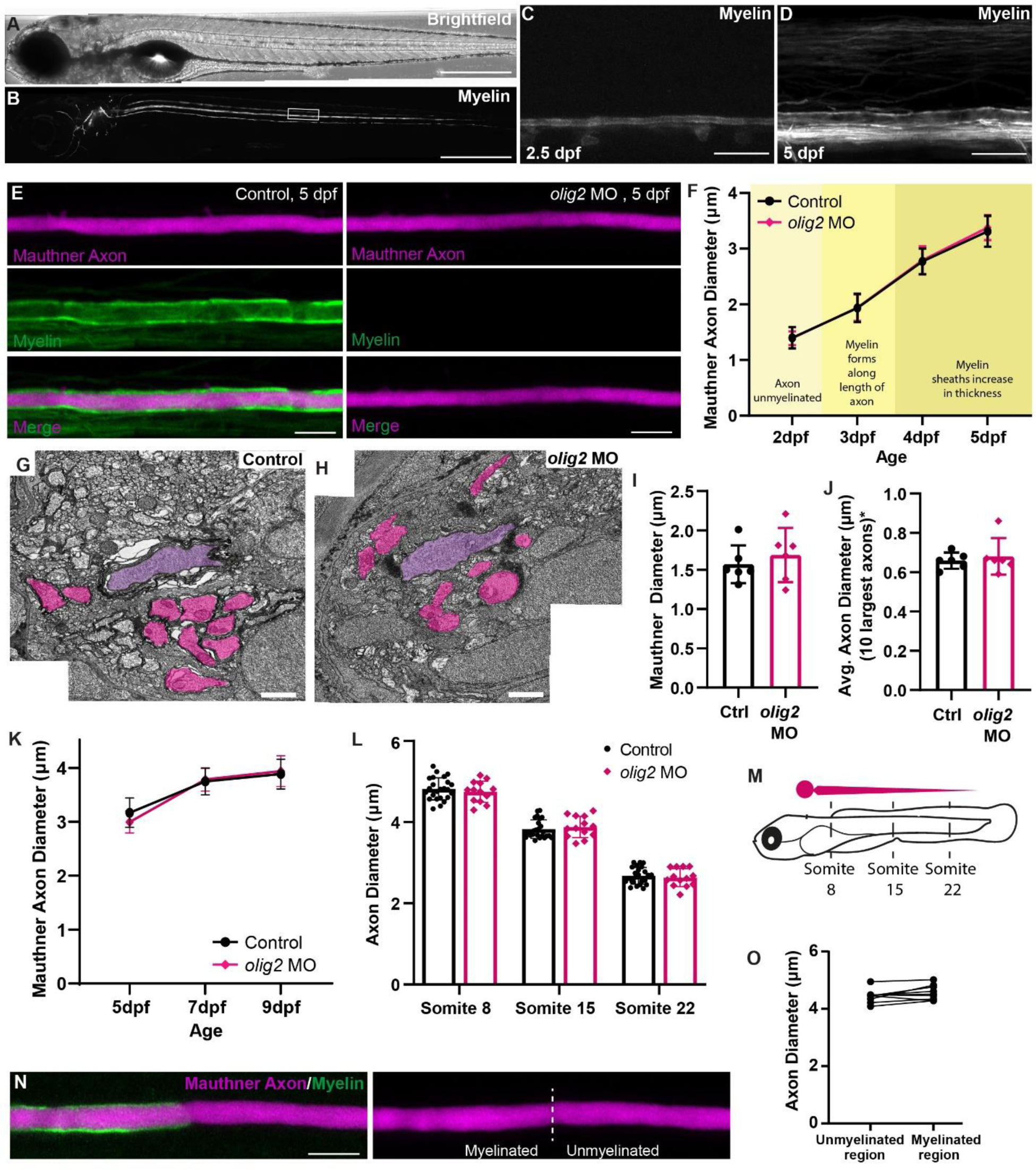
Absence of oligodendrocytes and myelin does not influence Mauthner axon diameter growth. (A) Brightfield images of 5 dpf larval zebrafish. (B) Fluorescent image of a 5 dpf larval zebrafish in which all myelin has been labelled using the transgenic reporter line mbp:EGFPCAAX. (C) Myelin in the zebrafish spinal cord (somite 15) at 2.5 dpf and (D) 5 dpf labelled using the transgenic reporter line mbp:EGFPCAAX. (E) Representative images of the Mauthner axon and oligodendrocytes/myelin at 5 dpf (somite 15) in control zebrafish and zebrafish injected with an *olig2* MO to eliminate oligodendrocytes and myelin. (F) Time course analysis of axon diameter growth (somite 15) for individual Mauthner axons imaged daily from 2 to 5 dpf. No significant difference between control and olig2 MO (n=13 control, 16 *olig2* MO axons from individual fish, 2-Way RM ANOVA, p=0.7864). (G) Representative transmission electron microscopy images of the 5 dpf ventral spinal cord (somite 15-16) in control and (H) *olig2* MO-injected zebrafish. Purple shading highlights the Mauthner axon and magenta shading highlights the next 10 largest axons (I) Mauthner axon diameter measured from 5 dpf transmission electron microscopy images (n=6 control, 6 *olig2* MO axons from individual fish, two-tailed unpaired t-test, p=0.5064). (J) Average diameter of the ten largest axons (*excluding Mauthner) in a hemi-ventral spinal cord (n=6 control, 6 *olig2* MO fish, two-tailed unpaired t-test, p=0.6124). (K) Time course analysis of axon diameter growth (somite 15) for individual Mauthner axons imaged from 5 to 9 dpf. No significant difference between control and *olig2* MO (n=20 control, 9 *olig2* MO axons from individual fish, 2-Way RM ANOVA, p=0.7693). (L) Mauthner axon diameter at 9dpf for the same axons at somite 8, 15, and 22 in control and *olig2* MO injected zebrafish. There are no significant differences between control and *olig2* MO (n=24 control, 13 *olig2* MO axons from individual fish, 2-Way RM ANOVA, p=0.7802). (M) Schematic depicting the locations imaged along the Mauthner axon in panel L. (N) Representative image of a partially myelinated Mauthner axon. (O) Comparison of axon diameter along adjacent unmyelinated and myelinated regions of a partially myelinated Mauthner axon at 7 dpf. There is no significant difference between myelinated and unmyelinated regions (n=9 axons from individual zebrafish, Two-tailed paired t-test, p=0.0826). Scale bars = 500 µm (A,B); 20 µm (C,D); 10 µm (E,N); 1 µm (G,H).

We first confirmed the absence of oligodendrocytes and myelin within the spinal cord by using the Tg(mbp:eGFP-CAAX) reporter line to fluorescently label oligodendrocytes (Figure 3 E), as well as by transmission electron microscopy (Figure 3 G,H). Next, we performed time-course live imaging and found that the absence of oligodendrocytes/myelin had no measurable effect on rapid diameter growth of the Mauthner axon that occurs during the time window in which it is usually myelinated, or the following days (Figure 3 F). In addition to the Mauthner axon, we also assessed the diameter of the next ten largest axons in the 5 dpf ventral spinal cord using electron microscopy (Figure 3 G,H). Similar to the Mauthner axon (Figure 3 I), no change in axon diameter was observed for these axons (Figure 3 J).

To determine whether myelination might have a delayed effect on diameter growth, we imaged the Mauthner axon at later time points, up to 9 dpf when its diameter growth begins to plateau. At this latest timepoint, we still did not observe any changes in Mauthner axon diameter in the absence of myelin (Figure 3 K). As distance from the cell body might potentially influence the reliance of the axon on myelin to support diameter growth, we extended our analyses at 9 dpf to include both proximal (somite 8) and distal (somite 22) portions of the axon, in addition to our standard measurements mid-way along the axon at somite 15 (Figure 3 M). While axon diameter did taper between the proximal to distal regions of the axon, no differences in axon diameter were observed between the control and *olig2* MO conditions (Figure 3 L).

In some *olig2* MO-treated larvae we did observe the occasional oligodendrocyte that did differentiate and form a myelin sheath along the Mauthner axon, usually along its proximal region. We took advantage of this to ask whether we could observe local changes along individual axons in response to the formation of myelin by comparing the diameter of the axon beneath the myelin to the diameter along the adjacent unmyelinated region (Figure 3 N). Again, we observed no significant differences in axon diameter along myelinated segments of sparsely myelinated Mauthner axons (Figure 3 O), further supporting the conclusion that interactions between myelin and the underlying axons do not locally modulate axon diameter.

## DISCUSSION

Together, our data from zebrafish and rodent models support the conclusion that the developmental growth of CNS axons to their normal and quite diverse diameters does not require ensheathment or myelination by oligodendrocytes. This contrasts with the PNS, where axons fail to reach appropriate diameters when they are not myelinated ^19, 21, 22^, highlighting a difference in how myelin produced by CNS oligodendrocytes and PNS Schwann cells affects axons. We also do not find a role for myelin in restricting axon diameter, as has been shown for CMTM6 expressed by Schwann cells in the PNS ^23^. Interestingly, the regulation of myelination by axon diameter also differs between the CNS and PNS. In the PNS, axon diameter is tightly linked to the amount of Neuregulin type III present on the axonal surface, which determines whether an axon is myelinated and the thickness of the myelin sheath, with axons over 1 μm typically being myelinated ^55, 56^. In contrast, no single axonal signal for myelination has been identified in the CNS, and myelination is less strictly correlated with diameter; axons between 0.2 μm and 1 μm may be either myelinated or unmyelinated ^57, 58, 59, 60^. The reasons for these differences in the reciprocal regulation of axon diameter and myelination between the CNS and PNS remain unclear, but may reflect distinct evolutionary pressures on conduction speed and space.

While we have shown that the developmental diameter growth of CNS axons does not depend on their myelination, it is noteworthy that previous studies indicate that axon diameters in the CNS can be affected by preventing the expression of certain myelin proteins^28, 30^ or following demyelination ^31, 61^. This highlights that while CNS myelin is not required for axons to achieve their appropriate and diverse diameters, its molecular, morphological or metabolic disruption can influence axonal size. Once myelinated and insulated from the extracellular environment, axon diameter growth and maintenance may, for example, become reliant on trophic and metabolic support from oligodendrocytes ^62, 63,64^. Interestingly, decreases to axon diameter following demyelination appear to be transient, as chronically demyelinated axons have been shown to recover their normal diameters over time ^31^, further supporting that it is the disruption of myelin, not its presence, that influences axon diameter.

It remains to be determined what are the primary factors regulating axon diameter within the CNS, and to what extent this is intrinsic to neuronal sub-type versus non-cell autonomously regulated. While our results argue against a major role for oligodendrocytes in axon diameter growth and diversity, the potential contribution of other glial cell types warrants further investigation. Given the importance of axon diameter in modulating conduction speeds, axon diameter growth may also be regulated by the neuronal circuit into which a neuron is integrated, for example, by levels of neuronal activity ^65, 66, 67^, signals from target cells ^18^, and/or number of synapses ^68^. Further studies will be essential to unravel the interplay between cell-intrinsic growth programmes and extrinsic regulatory cues that shape axon diameter. A deeper understanding of these mechanisms will provide critical insights into CNS development and the function of neuronal circuits.

## MATERIALS AND METHODS

### Mouse husbandry and transgenic lines

*Mbp^shi/shi^* ^33^ and littermate control mice were bred under a 12 hr light/dark cycle with food and water available ad libitum in the animal facility of the Max Planck Institute for Multidisciplinary Sciences (MPI-NAT, Göttingen, Germany) and tissue was dissected under license 33.19-42502-04-16/2337 issued by the Niedersächsisches Landesamt für Verbraucherschutz und Lebensmittelsicherheit (LAVES). The animal facility at the MPI-NAT is registered at the LAVES according to TierSchG §11 Abs. 1. According to the German Animal Welfare Law (Tierschutzgesetz der Bundesrepublik Deutschland, TierSchG) and the regulation about animals used in experiments, dated 11^th^ August 2021 (Tierschutz-Versuchstierverordnung, TierSchVersV), an animal welfare officer and an animal welfare committee are established for the institute.

Mice used to generate *Myrf* cKO mice were housed in a pathogen-free temperature-controlled environment on a 12 hr light/ dark cycle with food and water available *ad libitum* and animal procedures performed in accordance with, and approved by, the Oregon Health & Science Institutional Animal Care and Use Committee (TR02_IP00001328). *Myrf* conditional knock-out mice were generated by crossing *Myrf*^fl/fl^ mice ^46^ (B6;129-*Myrf^tm1Barr^* / J, JAX: 010607) to *Olig2*^wt/Cre^ mice ^69^ (Olig2^tm2(TVA,cre)Rth^ / J, JAX: 011103) for two generations. *Myrf*^fl/fl^; *Olig2*^wt/Cre^ mice were used as dysmyelinated mice with *Myrf*^wt/fl^; *Olig2*^wt/Cre^ littermates serving as aged-matched controls.

### *Mbp^shi/shi^* transmission electron microscopy and axon diameter analyses

*Mbp^shi/shi^* and littermate control optic nerves (P75) were immersion fixed in Karlsson–Schultz fixative (4% PFA, 2.5% glutaraldehyde in 0.1 M phosphate buffer) solution ^70^ for at least 24 h at 4°C and prepared using the reduced osmium tetroxide—thiocarbohydrazide—osmium (OTO) method as originally introduced by Deerinck and colleagues ^71^ with minor modifications as previously described ^28, 72, 73^. After resin polymerization, ultrathin sections (60–70 nm) were cut using an ultramicrotome (Leica UC-7, Vienna, Austria) and a 35° diamond knife (Diatome, Biel, Switzerland). The sections were placed on 100 mesh hexagonal copper grids (Science Services, Munich, Germany) and imaged with a Leo 912 electron microscope (Carl Zeiss, Oberkochen, Germany) and an on-axis 2 k CCD camera (TRS, Moorenweis, Germany). To determine axonal diameters, 10–15 random non-overlapping electron micrographs per mouse and 4–5 mice per genotype were analyzed. Analysis was performed using Fiji version 1.53c^74^. Per electron micrograph, up to 30 axons were selected at random using the Grid-Tool (Circular grids, 3 μm^2^ per point, random offset). The Feret diameter of all normal appearing axons comprised in the grid circles was analyzed.

### *Myrf* cKO transmission electron microscopy and axon diameter analyses

At P14 experimental mice were deeply anesthetized with a lethal dose of ketamine/xylazine (400mg/kg and 60mg/kg respectively) and subsequently perfused with 3 mL of phosphate buffered saline (PBS) followed by 15 mL of freshly hydrolyzed 4% paraformaldehyde (PFA, Electron Microscopy Sciences) in PBS. Optic nerves were dissected and processed for transmission electron microscopy largely as previously described^75^. Optic nerves were fixed in 2% PFA (15710, Electron Microscopy Sciences) and 2% Glutaraldehyde (16310, Electron Microscopy Sciences) in 0.1 M cacodylate buffer at room temperature for at least 2 hours, then moved into 4°C overnight. The following day, the nerves were stored in 1.5% PFA, 1.5% glutaraldehyde, 50 mM sucrose, 22.5 mM CaCl2•2H2O in 0.1 M cacodylate buffer for no more than one month. Optic nerves were then further fixed in 2% osmium-tetroxide (19190 Electron Microscopy Sciences) using a Biowave+ microwave (Ted Pella). The nerves went through subsequent acetone dehydration steps and embedding in Embed 812 (14120 Electron Microscopy Sciences). 400 nm semithin sections were cut using a Leica UC7 ultramicrotome, stained with 1% toluidine blue (Fisher Scientific) in 2% sodium tetraborate decahydrate (Fisher Scientific), and examined using a light microscope (Leica DM300) to ensure quality before cutting for TEM. Ultrathin sections (70 nm) were cut and placed on Formvar-coated copper grids (EMS), then counterstained with UranyLess (Electron Microscopy Sciences) and 3% lead citrate (Electron Microscopy Sciences). Images were acquired with an FEI Tecnai T12 TEM microscope equipped with an Advanced Microscopy Techniques (AMT) CCD camera.

To measure axon diameters, we used 17-22 non-overlapping electron micrographs (88.5 µm^2^ per image) sampled from across the optic nerve cross section per mouse from 4 mice per genotype. Analysis was performed using Fiji version 1.51n. For each image, axons were selected by placing a grid of crosses over each image using the Grid tool (crosses, 1.5 µm^2^, random offset). Due to differences in axon density between genotypes, ever second cross was selected for the *Myrf* cKOs to ensure a similar numbers of axons were sampled across all mice. Axons overlaid by crosses were traced using the polygon selection tool to obtain an area measurement, and the diameter calculated using the equation diameter = 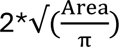.

### Zebrafish husbandry and transgenic lines

Adult zebrafish (*Danio rerio*) were housed and maintained in accordance with standard procedures in the Queen’s Medical Research Institute zebrafish facility at the University of Edinburgh, with both adult and larval zebrafish subject to a 14/10 hr, light/dark cycle. All experiments were performed in compliance with the UK Home Office, according to its regulations under project licenses 70/8436 and PP5258250.

The following stable, germline inserted transgenic lines were used in this study: Tg(mbp:eGFP-CAAX) ^47^, Tg(hspGFF62A:Gal4) ^76, 77^ and Tg(UAS:mRFP) ^76^. Embryos were produced by pairwise matings of homozygous Tg(mbp:eGFP-CAAX) x homozygous Tg(hspGFF62A:Gal4); Tg(UAS:mRFP) zebrafish. Embryos were raised at 28.5 C°, with 50 embryos (or less) per 10 cm petri dish in ∼45 mL of 10 mM HEPES-buffered E3 embryo medium or conditioned aquarium water with methylene blue. Larvae were staged according to days post-fertilisation (dpf), with larvae between 2 dpf and 9 dpf used in this study. At these ages, sexual differentiation of zebrafish has not yet occurred.

### Eliminating myelin using *olig2* MO

In order to inhibit oligodendrocyte development in the zebrafish embryo, 7.5 ng of *olig2* morpholino (Gene Tools, LLC, Philomath, USA) in nuclease-free water was injected into fertilized eggs at 1-4 cell stages. Olig2-ATG-MO sequence: 5’-ACACTCGGCTCGTGTCAGAGTCCAT 3’ ^54^.

### Live-imaging and quantification of axon diameter

Zebrafish larvae were anaesthetised with 600 µM tricaine in E3 embryo medium and immobilised in 1.3% low melting-point agarose on a glass coverslip, which was suspended over a microscope slide using high vacuum silicone grease to create a well containing E3 embryo medium and 600 µM tricaine. Z-stacks (with optimal z-step) were obtained using a Zeiss LSM880 microscope with Airyscan FAST in super-resolution mode, using a Zeiss Plan-Apochromat 20x dry, NA = 0.8 objective, and processed using the default Airyscan processing settings (Zen Black 2.3, Zeiss). Images of the Mauthner axon and myelin were taken from a lateral view of the spinal cord centred around somite 15, unless otherwise indicated.

Axon diameters from AiryScan FAST confocal images were measured in Fiji using custom macros (available at https://github.com/jasonjearly/Axon_Caliber/releases/tag/v1.0.0) as previous described ^48^. Briefly, a “Split Axons Tool” was used on the whole spinal cord z-stack datasets to separate the two Mauthner axons and generate a maximum projection image of each axon. The maximum intensity projection for the axon closest located closest to the imaging objective was then used to measure axon diameter. To measure axon diameter, an “Axon Trace Tool” was used to trace the centre point of the axon along its length. Then the “Axon Caliber Tool” was used to measure the average axon diameter along the length of the axon selected by the Axon Trace Tool.

### Zebrafish transmission electron microscopy

Zebrafish tissue was prepared for transmission electron microscopy according to a previously published protocol ^78^. Briefly, zebrafish embryos were terminally anaesthetised in tricaine and incubated with primary fixative of 4% paraformaldehyde + 2% glutaraldehyde in 0.1 M sodium cacodylate buffer with microwave stimulation (100 W for 1 min ON, 1 min OFF, 1 min ON followed by 450 W for 20 s ON, 20s OFF repeated five times), followed by 3 hr incubation at room temperature. Tissue was then stored for 5 days at 4°C in a 2% glutaraldehyde post-fixative. Samples were washed in 0.1 M Cacodylate buffer and incubated with microwave stimulation (100 W for 1 min ON, 1 min OFF, 1 min ON followed by 450 W for 20 s ON, 20s OFF repeated five times) in secondary fixative of 2% osmium tetroxide in 0.1 M sodium cacodylate/0.1 M imidazole buffer pH7.5, then left 3 hours at room temperature. Samples were washed with distilled water, then stained en bloc with a saturated (8%) uranyl acetate solution with microwave stimulation (450 W for 1 min ON, 1 min OFF, 1 min ON) followed by overnight incubation at room temperature. Next, samples were dehydrated with an ethanol series and acetone using microwave stimulation (250 W for 45s). Samples were embedded in Embed-812 resin (Electron Microscopy Sciences) and sections (70-80 nm in thickness, silver) cut using a Leica Ultracut Microtome and diamond knife (Diatome). Sections were mounted on hexagonal copper electron microscopy grids (200 Mesh Grids, Agar Scientific). Mounted sections were stained in uranyl acetate and Sato’s lead stain (refer to ^78^ for stain preparation). TEM imaging was performed at the University of Edinburgh Biology Scanning Electron Microscope Facility using a Jeol JEM1400 Plus Transmission Electron Microscope. The ventral spinal cord was imaged at 12000x magnification. In order to create panoramic views, individual electron micrograph tiles were aligned using the automated photomerge tool in Adobe Photoshop 2020 (v21.0.2). To assess axon diameter, the Mauthner axon, as well as the next 10 largest axons in the ventral spinal cord were traced in Fiji (v1.51n) to obtain cross-sectional areas, which were then used to calculate diameter using the equation diameter = 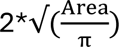.

### Statistical analysis

Statistical analysis was performed using GraphPad Prism 10 (up to version 10.2.3) and R version 4.4.1 (The R Foundation for Statistical Computing), with results included in the figure legends. Significance was defined as p < 0.05. For evaluation of differences between axon diameter means for mouse datasets, the data were fit with linear mixed effect models using the R package lme4^79^, and comparisons made using ANOVA to compare the model containing genotype information with a null model. Evaluation of axon diameter distributions for mouse data sets was carried out using the package Linear Quantile Mixed Models (lqmm)^80^, with Nelder-Mead optimization and maximum iterations set to 5000. Estimation of p values for the coefficients for each quantile were obtained using the block-bootstrap method with 500 iterations.

## ACKNOWLEDGEMENTS

We thank the University of Edinburgh BVS Zebrafish Facility, Zebrafish Imaging and Screening Facility, and Transmission Electron Microscopy Facility for expert assistance, and Dr. L. de Hoz for breeding and supplying shiverer mice. This work was supported by Wellcome Trust Senior Research Fellowships (102836/Z/13/Z and 214244/Z/18/Z) and a Lister Institute Research Prize to D.A.L.. J.M.B. was supported by a Canadian Institute of Health Research a postdoctoral fellowship and a Multiple Sclerosis Society of Canada/ Fonds de la recherche du Quebec-Sante postdoctoral fellowship. R.A.D. and K.R.M. were supported by R01NS120651. B.E. and work in his laboratory were supported by NIH award R01NS120981 and an endowment from the Warren family. K.E. was supported by NINDS T32AG055378.

---

**Supplemental Figure 1:**
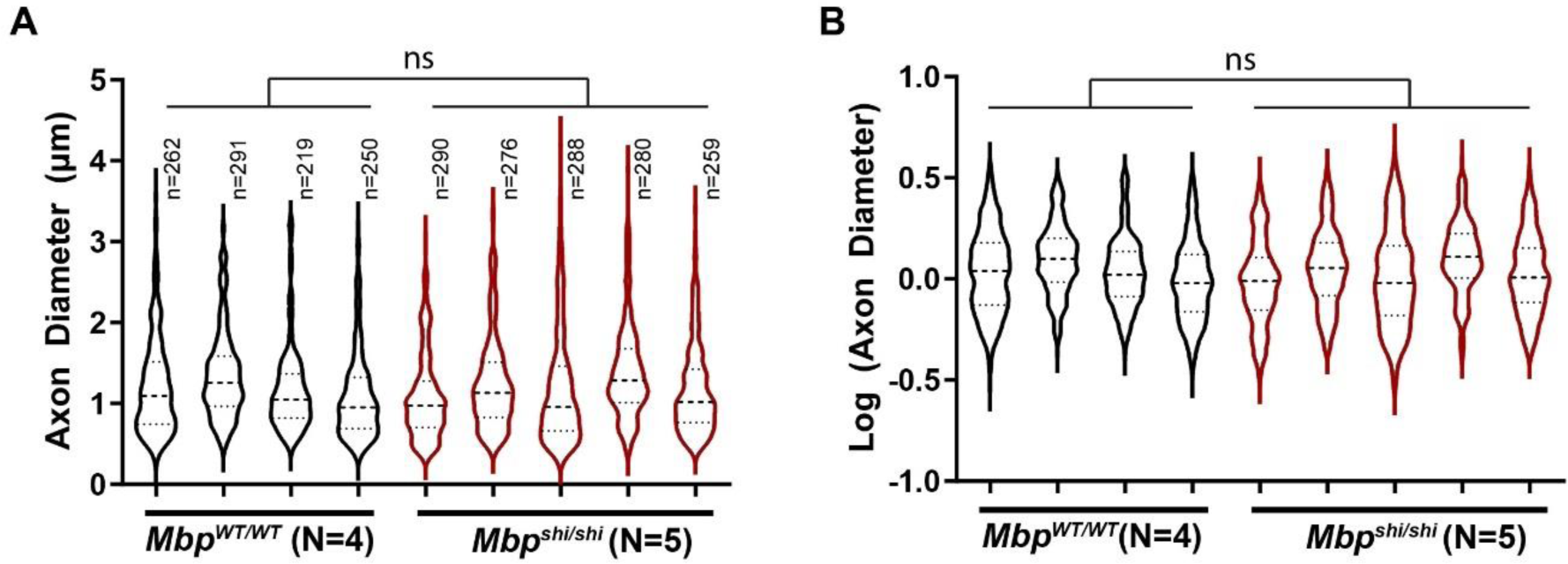
Normal axon diameters in *Mbp^shi/shi^* optic nerves at P75. (A) Axon diameter distribution for each animal. Solid horizontal line marks the median and horizontal dotted lines mark the quartiles. To evaluate possible differences between groups, while also accounting for all observations made from each animal, we fit the data with linear mixed effects models that included animal identity as a random effect and genotype as a fixed effect. Comparison to a null model did not reveal any significant effect of genotype on the mean diameter (p = 0.939, Chisq = 0.0058, df = 1, chi squared test). Evaluation of the distribution of the diameters using quantile analysis did not identify differences at the quartile boundaries (first and second quartile boundary: p = 0.74; median: p = 0.74; third and fourth quartile boundary: p = 0.70, block-bootstrap). (B) Log transformation of data in (A). A similar analysis to (A) also did not reveal a significant effect of genotype on the mean axon diameter (p = 0.836, Chisq = 0.0429, df = 1) or differences at the quartile boundaries (first and second quartile boundary: p = 0.68; median: p = 0.75; third and fourth quartile boundary: p = 0.71, block-bootstrap).

**Supplemental Figure 2:**
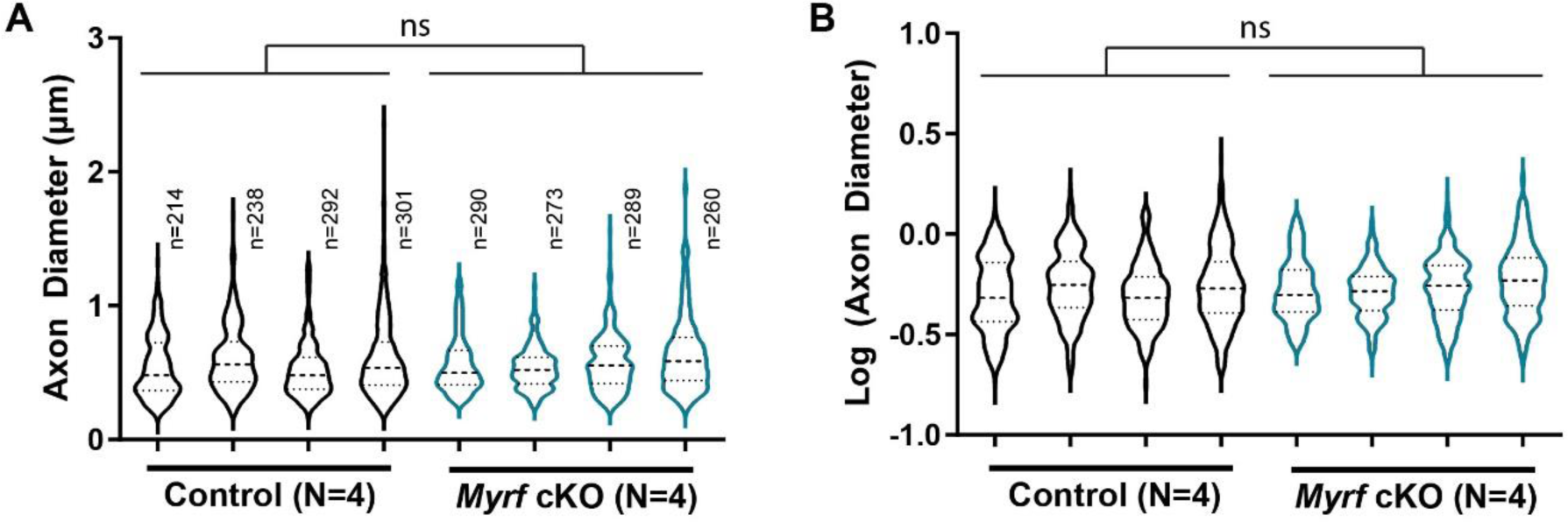
Normal axon diameters in *Myrf* cKO optic nerves at P14. (A) Axon diameter distribution for each animal. Solid horizontal line marks the median and horizontal dotted lines mark the quartiles. Evaluation of linear mixed effects models that included animal identity as a random effect and genotype as a fixed effect did not reveal any significant effect of genotype on the mean diameter (p = 0.80, Chisq = 0.0664, df = 1, chi squared test). Evaluation of the distribution of the diameters using quantile analysis suggested a possible defect of genotype on the boundary between the first and second quartiles (p = 0.0046, block bootstrap) did not identify differences in the median or at the boundary between the third and fourth quartile (median: p = 0.88; third and fourth quartile boundary: p = 0.80). (B) Log transformation of data in (A). A similar analysis to (A) also did not reveal a significant effect of genotype on the mean axon diameter (p = 0.406, Chisq = 0.691, df = 1) or differences at the quartile boundaries (first and second quartile boundary: p = 0.086; median: p = 0.40; third and fourth quartile boundary: p = 0.79, block-bootstrap).

