## Supplemental Figures for "Developmental axon diameter growth of central nervous system axons does not depend on ensheathment or myelination by oligodendrocytes"

Supplemental Figures 1 and 2

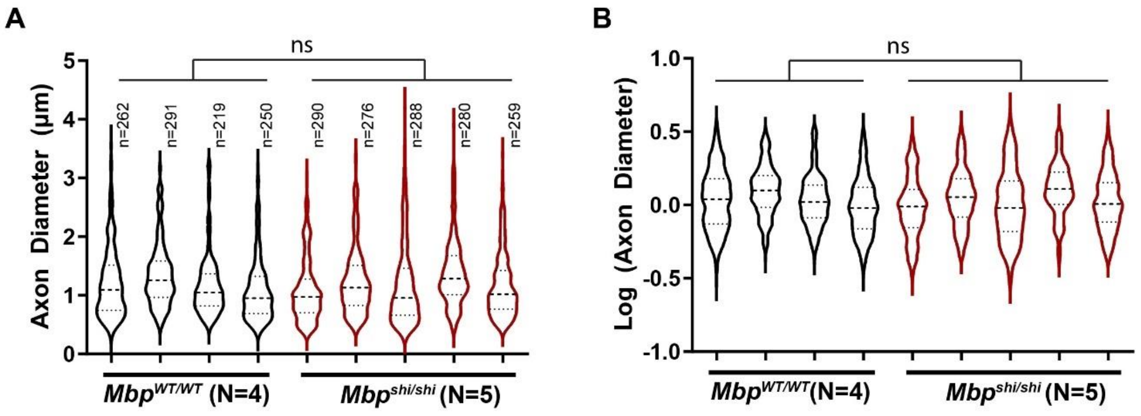


**Supplemental Figure 1: Normal axon diameters in *Mbp*$^{shi/shi}$ optic nerves at P75.** (A) Axon diameter distribution for each animal. Solid horizontal line marks the median and horizontal dotted lines mark the quartiles. To evaluate possible differences between groups, while also accounting for all observations made from each animal, we fit the data with linear mixed effects models that included animal identity as a random effect and genotype as a fixed effect. Comparison to a null model did not reveal any significant effect of genotype on the mean diameter (p = 0.939, Chisq = 0.0058, df = 1, chi squared test). Evaluation of the distribution of the diameters using quantile analysis did not identify differences at the quartile boundaries (first and second quartile boundary: p = 0.74; median: p = 0.74; third and fourth quartile boundary: p = 0.70, block-bootstrap). (B) Log transformation of data in (A). A similar analysis to (A) also did not reveal a significant effect of genotype on the mean axon diameter (p = 0.836, Chisq = 0.0429, df = 1) or differences at the quartile boundaries (first and second quartile boundary: p = 0.68; median: p = 0.75; third and fourth quartile boundary: p = 0.71, block-bootstrap).

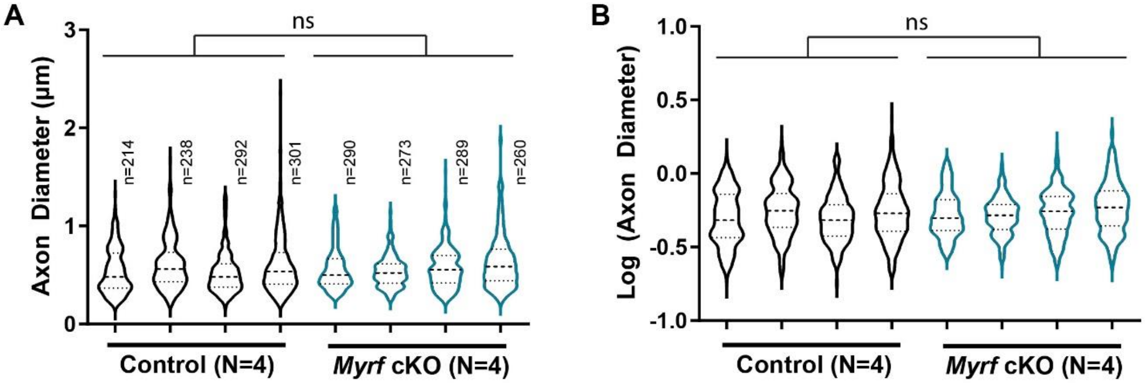


**Supplemental Figure 2: Normal axon diameters in *Myrf* cKO optic nerves at P14.** (A) Axon diameter distribution for each animal. Solid horizontal line marks the median and horizontal dotted lines mark the quartiles. Evaluation of linear mixed effects models that included animal identity as a random effect and genotype as a fixed effect did not reveal any significant effect of genotype on the mean diameter (p = 0.80, Chisq = 0.0664, df = 1, chi squared test). Evaluation of the distribution of the diameters using quantile analysis suggested a possible defect of genotype on the boundary between the first and second quartiles (p = 0.0046, block bootstrap) did not identify differences in the median or at the boundary between the third and fourth quartile (median: p = 0.88; third and fourth quartile boundary: p = 0.80). (B) Log transformation of data in (A). A similar analysis to (A) also did not reveal a significant effect of genotype on the mean axon diameter (p = 0.406, Chisq = 0.691, df = 1) or differences at the quartile boundaries (first and second quartile boundary: p = 0.086; median: p = 0.40; third and fourth quartile boundary: p = 0.79, block-bootstrap).
